# Imputation of Spatially-resolved Transcriptomes by Graph-regularized Tensor Completion

**DOI:** 10.1101/2020.08.05.237560

**Authors:** Zhuliu Li, Tianci Song, Jeongsik Yong, Rui Kuang

## Abstract

High-throughput spatial-transcriptomics RNA sequencing (sptRNA-seq) based on in-situ capturing technologies has recently been developed to spatially resolve transcriptome-wide mRNA expressions mapped to the captured locations in a tissue sample. One major limitation of in-situ capturing is the high dropout rate of mRNAs that fail the capture or the amplification, which leads to incomplete profiling of the gene expressions. In this paper, we introduce a graph-regularized tensor completion model for imputing the missing mRNA expressions in sptRNA-seq data, namely FIST, Fast Imputation of Spatially-resolved transcriptomes by graph-regularized Tensor completion. We first model sptRNA-seq data as a 3-way sparse tensor in genes (*p*-mode) and the (*x, y*) spatial coordinates (*x*-mode and *y*-mode) of the observed gene expressions, and then consider the imputation of the unobserved entries as a tensor completion problem in Canonical Polyadic Decomposition (CPD) form. To improve the imputation of highly sparse sptRNA-seq data, we also introduce a protein-protein interaction network to add prior knowledge of gene functions, and a spatial graph to capture the the spatial relations among the capture spots. The tensor completion model is then regularized by a Cartesian product graph of protein-protein interaction network and the spatial graph to capture the high-order relations in the tensor. In the experiments, FIST was tested on ten 10x Genomics Visium spatial transcriptomic datasets of different tissue sections with cross-validation among the known entries in the imputation. FIST significantly outperformed several best performing single-cell RNAseq data imputation methods. We also demonstrate that both the spatial graph and PPI network play an important role in improving the imputation. In a case study, we further analyzed the gene clusters obtained from the imputed gene expressions to show that the imputations by FIST indeed capture the spatial characteristics in the gene expressions and reveal functions that are highly relevant to three different kinds of tissues in mouse kidney. The source code and data are available at https://github.com/kuanglab/FIST.

**Author summary:** Biological tissues are composed of different types of structurally organized cell units playing distinct functional roles. The exciting new spatial gene expression profiling methods have enabled the analysis of spatially resolved transcriptomes to understand the spatial and functional characteristics of these cells in the context of eco-environment of tissue. Similar to single-cell RNA sequencing data, spatial transcriptomics data also suffers from a high dropout rate of mRNAs in in-situ capture. Our method, FIST (Fast Imputation of Spatially-resolved transcriptomes by graph-regularized Tensor completion), focuses on the spatial and high-sparsity nature of spatial transcriptomics data by modeling the data as a 3-way gene-by-(*x, y*)-location tensor and a product graph of a spatial graph and a protein-protein interaction network. Our comprehensive evaluation of FIST on ten 10x Genomics Visium spatial genomics datasets and comparison with the methods for single-cell RNA sequencing data imputation demonstrate that FIST is a better method more suitable for spatial gene expression imputation. Overall, we found FIST a useful new method for analyzing spatially resolved gene expressions based on novel modeling of spatial and functional information.

## Introduction

Dissection of complex genomic architectures of heterogeneous cells and how they are organized spatially in tissue are essential for understanding the molecular and cellular mechanisms underlying important phenotypes. For example, each tumor is a mixture of different types of proliferating cancerous cells with changing genetic materials [1]. The cancer cell sub-populations co-evolve in the micro-environment formed around their spatial locations. It is important to understand the cell-cell interactions and signaling as well as the functioning of each individual cell to develop effective cancer treatment to eradicate all cancer clones at their locations [2]. Conventional gene expression analyses have been limited to low-resolution bulk profiling that measures the average transcription levels in a population of cells. With single-cell RNA sequencing (scRNA-seq) [3–5], single cells are isolated with a capture method such as fluorescence-activated cell sorting (FACS), Fluidigm C1 or microdroplet microfluidics and then the RNAs are captured, reverse transcribed and amplified for sequencing the RNAs barcoded for the individual origin cells [6,7]. While scRNA-seq is useful for detecting the cell heterogeneity in a tissue sample, it does not provide the spatial information of the isolated cells. To map cell localization, earlier in-situ hybridization methods such as FISH [8], FISSEQ [9], smFISH [10] and MERFISH [11] were developed to profile up to a thousand targeted genes in pre-constructed references with single-molecule RNA imaging. Based on in-situ capturing technologies, more recent spatial transcriptomics RNA sequencing (sptRNA-seq) [12–15] combines positional barcoded arrays and RNA sequencing with single-cell imaging to spatially resolve RNA expressions in each measured spot in the array [12,16–18]. These new technologies have transformed the transcriptome analysis into a new paradigm for connecting single-cell molecular profiling to tissue micro-environment and the dynamics of a tissue region [19–21].

With in-situ capturing technology, mRNAs are captured and sequenced in the spots on the spatial genomic array aligned to the locations on the tissue. For example, spatial transcriptome techniques based on 10x Genomics Visium kit report the counts of mRNAs by unique molecular identifiers (UMIs) in the read-pairs mapped to each gene [22]. Very similar to scRNA-seq data, a significant technical limitation of sptRNA-seq data is known as *dropout*, where dropout events refer to the false quantification of a gene as unexpressed due to the failure in amplifying the transcripts during reverse-transcription [23]. For example, spatial transcriptomic technology’s detection efficiency is as low as 6.9% while 10x Genomics Visum has a slightly higher efficiency [24]. It has been shown in previous studies on scRNA-seq data that normalizations will not address the dropout effects [22, 25]. In the literature, many imputation methods such as Zero-inflated factor analysis (ZIFA) [26], Zero-Inflated Negative Binomial-based Wanted Variation Extraction (ZINB-WaVE) [27] and BISCUIT [25] have been developed to impute scRNA-seq. While these methods are also applicable to impute the spatial gene expressions, they ignore a unique characteristic of sptRNA-seq data, which is the spatial information among the gene expressions in the spatial array, and do not fully take advantage of the functional relations among genes for more reliable joint imputation.

To provide a more suitable method for imputation of spatially-resolved gene expressions, we introduce FIST, Fast Imputation of Spatially-resolved transcriptomes by graph-regularized Tensor completion. FIST is a tensor completion model regularized by a product graph as illustrated in Figure 1. FIST models sptRNA-seq data as a 3-way sparse tensor in genes (*p*-mode) and the (*x, y*) spatial coordinates (*x*-mode and *y*-mode) of the observed gene expressions (Figure 1(A)). As shown in Figure 1(B), a protein-protein interaction network models the interactions between pairs of genes in the gene mode, and the spatial graph is modeled by a product graph of two chain graph for columns (*x*-mode) and rows (*y*-mode) in the grid to capture the spatial relations among the (*x, y*) spots. The tensor product of the two graphs with prior knowledge of gene functions and the spatial relations among the capture spots are then introduced as a regularization of tensor completion to obtain the Canonical Polyadic Decomposition (CPD) of the tensor. The imputation of the unobserved entries can then be derived by reconstructing the entries in the completed tensor shown in Figure 1(C). In the experiments, we comprehensively evaluated FIST on ten 10x Genomics Visium spatial genomics datasets by comparison with widely used methods for single-cell RNA sequencing data imputation. We also analyzed a mouse kidney dataset with more functional interpretation of the gene clusters obtained by the imputed gene expressions to detect highly relevant functions in the clusters expressed in three kidney tissue regions, corex, outer stripe of the outer medulla (OSOM) and inner stripe of the outer medulla (ISOM).

**Fig 1.**
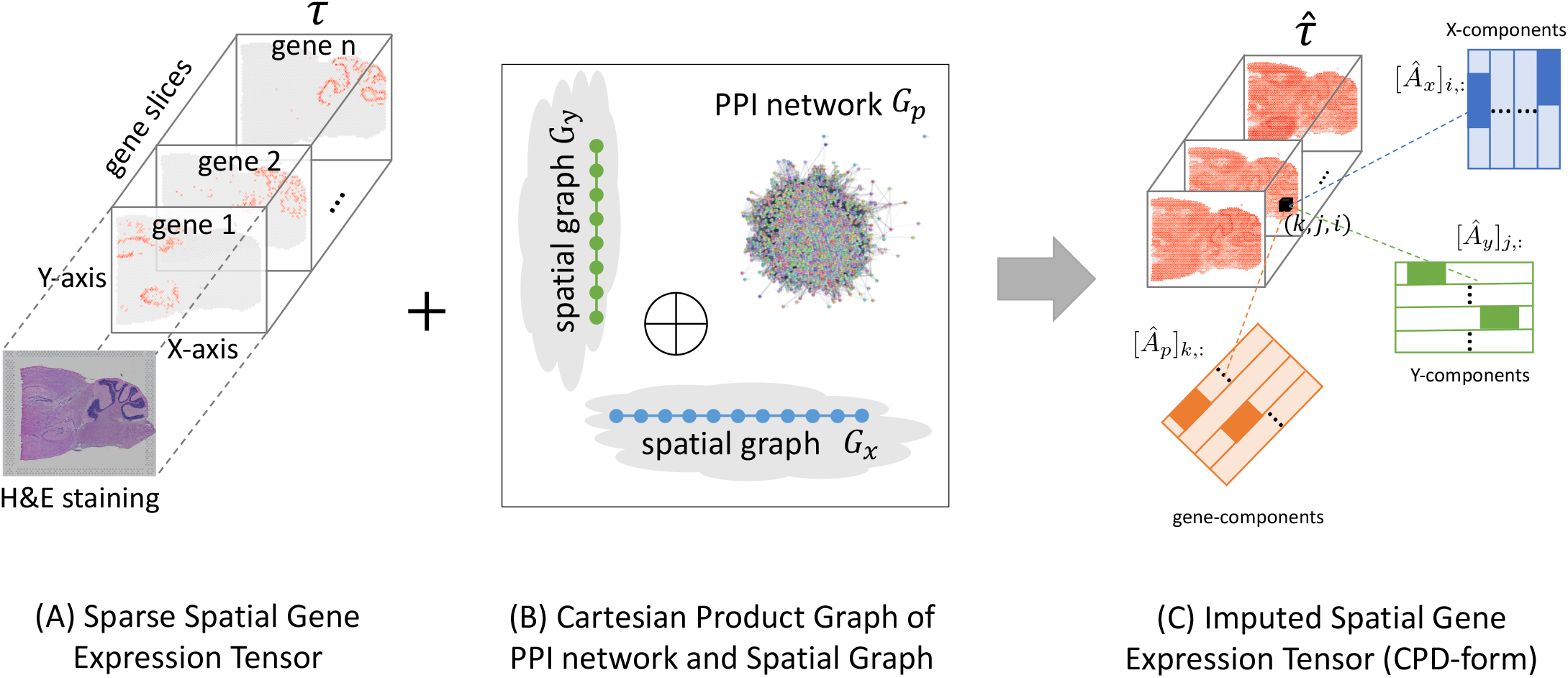
Imputation of spatial transcriptomes by graph-regularized tensor completion. **(A)** The input sptRNA-seq data is modeled by a 3-way sparse tensor in genes (*p*-mode) and the (*x, y*) spatial coordinates (*x*-mode and *y*-mode) of the observed gene expressions. H&E image is also shown to visualize the cell morphologies aligned to the spots. **(B)** A protein-protein interaction network and a spatial graph are integrated as a product graph for tensor completion. The spatial graph is also a product graph of two chain graph for columns (x-mode) and rows (y-mode) in the grid. **(C)** After the imputation, tire CPD form of the complete tensor can be used to impute any missing gene expressions, e.g. the entry (*k, j, i*) can be reconstructed as the sum of the element-wise multiplications of the three components [*Â_p_*]_*k*,:_, [*Â_y_*]_*j*,:_ and [*Â_x_*]_*i*,:_.

## Materials and methods

In this section, we first describe the task of spatial gene expression imputation, and next introduce the mathematical model for graph-regularized tensor completion problem. We then present a fast iterative algorithm FIST to solve the optimization problem defined to optimize the model. We also provide the convergence analysis of proposed algorithm in Appendix. Finally, we provide a review of several state-of-the-art methods for scRNA-seq data imputation, which are also compared in the experiments later. The notations which will be used for the derivations in the forthcoming sections are summarized in Table 1.

**Table 1.** Notations

| Notation | Definition |
| --- | --- |
| $G_x, G_y$ | Spatial graph of $(x, y)$ coordinates |
| $G_p$ | Protein-protein interaction (PPI) network |
| $n_x, n_y, n_p$ | Number of vertices in $G_x, G_y$ and $G_p$ |
| $W_x \in \mathbb{R}_{[0,1]}^{n_x \times n_x}, W_y \in \mathbb{R}_{[0,1]}^{n_y \times n_y}, W_p \in \mathbb{R}_{[0,1]}^{n_p \times n_p}$ | Adjacency matrix of $G_x, G_y, G_p$ |
| $L_x \in \mathbb{R}^{n_x \times n_x}, L_y \in \mathbb{R}^{n_y \times n_y}, L_p \in \mathbb{R}^{n_p \times n_p}$ | Graph Laplacian of $G_x, G_y, G_p$ |
| $\mathfrak{G}(x, y, z)$ | Cartesian product of $G_x, G_y$ and $G_z$ |
| $\mathfrak{W}(x, y, z) \in \mathbb{R}_{[0,1]}^{n_x n_y n_p \times n_x n_y n_p}$ | Adjacency matrix of $\mathfrak{G}(x, y, z)$ |
| $\mathfrak{L}(x, y, z) \in \mathbb{R}^{n_x n_y n_p \times n_x n_y n_p}$ | Graph Laplacian of $\mathfrak{G}(x, y, z)$ |
| $\mathcal{T} \in \mathbb{R}_+^{n_p \times n_y \times n_x}$ | Incomplete spatial gene expression tensor |
| $\hat{\mathcal{T}} \in \mathbb{R}_+^{n_p \times n_y \times n_x}$ | Complete spatial gene expression tensor |
| $\mathcal{M} \in \mathbb{R}_{[0,1]}^{n_p \times n_y \times n_x}$ | Binary mask tensor |
| $\hat{A}_x \in \mathbb{R}_+^{n_x \times r}, \hat{A}_y \in \mathbb{R}_+^{n_y \times r}, \hat{A}_p \in \mathbb{R}_+^{n_p \times r}$ | CPD component matrices of $\hat{\mathcal{T}}$ |
| $\text{vec}(\mathcal{T}) \in \mathbb{R}^{n_x n_y n_p \times 1}$ | Convert $\mathcal{T}$ to be a vector |

### Imputation of spatial gene expressions by tensor modeling

Let 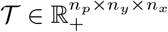 be the 3-way sparse tensor of the observed spatial gene expression data as show in Figure 1(A), with its zero entries representing the missing gene expressions, where *n_p_* denote the total number of genes, *n_x_* and *n_y_* denote the dimensions of the *x* and *y* spatial coordinates of the spatial transcriptomics array. Our goal is to learn a complete spatial gene expression tensor 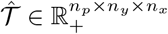 from 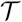 as illustrated in Figure 1(C). Apparently, it becomes computationally expensive and often infeasible to compute or store a dense tensor 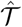, especially in high spatial resolutions with millions of spots. Therefore, we propose to compute an economy-size representation of 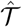 via an equality constraint 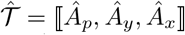, which is called *Canonical Polyadic Decomposition (CPD)* [28] of 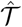 defined below

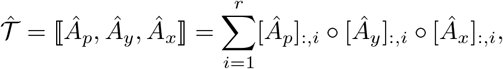

where *r* is the rank of 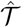, and o denotes the vector outer product. Here, [*Ã_x_*]_:*i*_ is the *i*-th column of the low-rank matrix 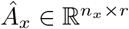, which can be similarly defined for [*Â_y_*]_:*i*_ and [*Â_p_*]_:*i*_. By utilizing the tensor CPD form, we replaced the optimization variables from 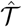 to *Â_p_, Â_y_* and *Â_x_*, reducing the number of parameters from *n_p_n_y_n_x_* to *r*(*n_p_* + *n_y_* + *n_x_*). The advantage of the tensor representation is to incorporate the 2-D spatial *x*-mode and *y*-mode such that the grid structure is preserved within the columns and the rows of the spatial array in the tensor, which contains useful spatial information. Next, we introduce the tensor completion model over *Â_p_*, *Â_y_* and *Â_x_*.

### Graph regularized tensor completion model

The key ideas of modeling the task of spatial gene expression imputation are i) the inferred complete spatial gene expression tensor 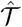 is regularized to integrate the spatial arrangements of the spot in the tissue array and the functional relations among the genes; ii) the observed part in 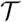 is also required to be preserved in 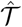 as the completion task requires; and iii) the inferred tensor 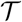 is compressed as the economy-size representation 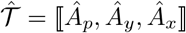 for scalable space and time efficiencies. The novel optimization formulation is shown below in Proposition 1,

#### Proposition 1.

*The complete spatial gene expression tensor* 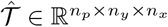 *can be obtained by solving the following optimization problem:*

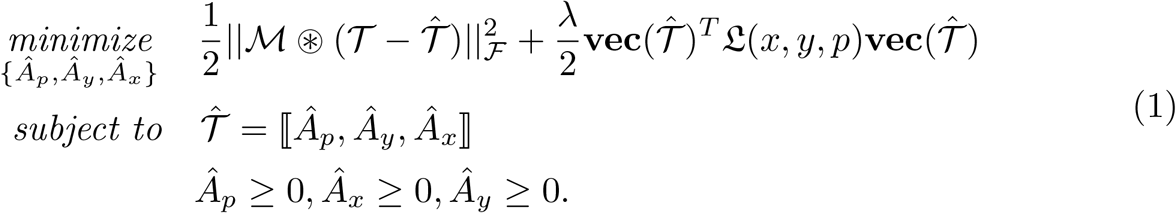

*where λ* ∈ [0,1] *is a model hyperparameter;* 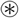 *denotes the Hadamard product; and* 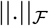 *denotes the Frobenius norm of a tensor*.

There are two optimization terms in the model defined in equation (1), consistency with the observations (the first term) and Cartesian product graph regularization (the second term), which are explained below,

- **Consistency with the observations** We introduce a binary mask tensor 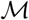 to indicate the indices of the observed entries in 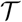. The (*i, j, k*)-th entry 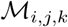 which is defined below, represents whether the (*i, j, k*)-th element in 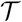 is observed or not.

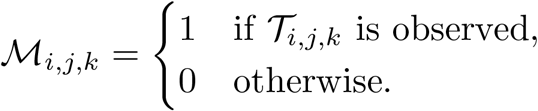 By introducing the squared-error in 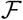-norm 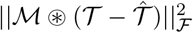 in the model, we ensure the inferred spatial gene expression tensor 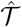 is consistent with its observed counterparts in 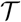.
- **Cartesian product graph regularization** Two useful assumptions to introduce prior knowledge for inferring the tensor are *1) the spatially adjacent spots should share similar gene expressions, and 2) the expression of two genes are likely highly correlated if they share similar gene functions [29, 30]*. We introduce a spatial graph and a protein-protein interaction (PPI) network into the model. We first encode the spatial information in two undirected unweighted graph *G_x_* = (*V_x_, E_x_*) and *G_y_* = (*V_y_, E_y_*), where *V_x_* and *V_y_* are vertex sets and *E_x_* and *E_y_* are edge sets. There are [*V_x_*] = *n_x_* vertices in *G_x_* where *n_x_* is the number of the spatial coordinates along the *x*-axis of the spatial array. Two vertices in *G_x_* are connected by an edge if they are adjacent along the *x*-axis. The connections in *G_y_* can be similarly defined to encode the *y*-coordinates of the tissue. We also incorporate the topological information of a PPI network download from BioGRID 3.5 [31] to use the functional modules in the PPI network. We denote the PPI network as *G_p_* = (*V_p_, E_p_*) which contains |*E_p_*| experimentally documented physical interactions among the |*V_p_*| = *n_p_* proteins. We then use the Cartesian product [32] 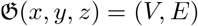 of the three individual graphs *G_x_, G_y_* and *G_p_* to regularize the elements in 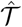, where |*V*| = *n_x_n_y_n_p_*. The (*v_x_v_y_v_p_*)-th vertex in *V* represents a triple of vertices {*v_x_* ∈ *V_x_, v_y_* ∈ *V_y_, v_p_* ∈ *V_p_*} from each of the three graphs. The (*a_x_, a_y_, a_p_*)-th and (*b_x_, b_y_, b_p_*)-th vertices in *V* are connected by an edge iff for any *i,j* ∈ {*x, y, p*}, (*a_i_, b_i_*) ∈ *E_i_* and *a_j_* = *b_j_* ∈ *V_j_* for all *j* ≠ *i*. Denoting the adjacency, degree and graph Laplacian matrices of graph *G_i_* as *W_i_, D_i_* and *L_i_* = *D_i_* – *W_i_* for *i* ∈ {*x, y, p*}, the adjacency and graph Laplacian matrices of 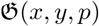 are obtained as 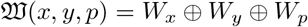 and 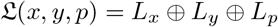 respectively, where ⊕ denotes the Kronecker sum [33]. By introducing the term 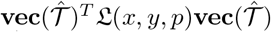 in equation (1), the inferred gene expression values in 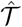 are ensured to be smooth over the manifolds of the product graph 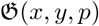, such that a pair of tensor entries 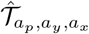 and 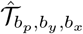 share similar values if the (*a_x_, a_y_, a_p_*)-th and (*b_x_, b_y_, b_p_*)-th vertices are connected in 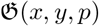. A connection implies that the *x*-coordinate *a_x_* and *b_x_* is adjacent or *y*-coordinate *a_y_* and *b_y_* is adjacent or gene *a_p_* and gene *b_p_* are connected in the PPI, with the two other dimensions fixed. Using Cartesian product graph is a more conservative strategy to connect multi-relations in a high-order graph as we have shown in [34] since only replacing one of the dimensions by the immediate neighbors is allowed to create connections. Note that it also possible to use tensor product graph or strong product graph [34] but there could be too many connections to provide meaningful connectivity in the product graph for helpful regularization.

### FIST Algorithm

The model introduced in equation (1) is non-convex on variables {*Â_p_, Â_y_, Â_x_*} jointly, thus finding its global minimum is difficult. In this section, we propose an efficient iterative algorithm **F**ast **I**mputation of **Sp**atially-resolved Gene Expression **T**ensor (FIST) to find its local optimal solution using the multiplicative updating rule [35], based on derivatives of *Â_p_, Â_y_* and *Â_x_*. Without loss of generality, we only show the derivations with respect to *Â_p_*, and provide the FIST algorithm in Algorithm 1.

We first bring the equality constraint 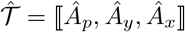 in Model (1) into the objective function, and rewrite the objective function as

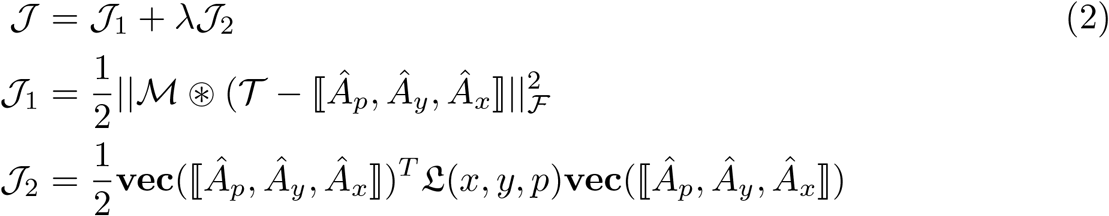

The partial derivative of 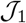 with respect to *Â_p_* can be computed as

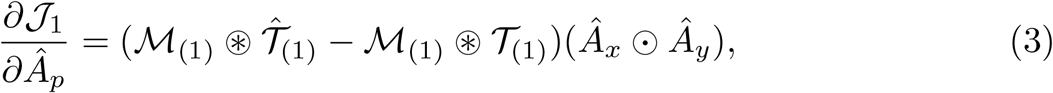

where 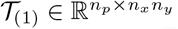 denotes the matrix flattened from tensor 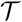; ⊙ denotes the Khatri–Rao product [28]. Note that the term 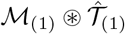 in Equation (3) implies we only need to compute the entries in 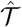 which correspond to the non-zero entries (indices of the observed gene expression) in 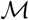. The rest of the computation in Equation (3) involves the well-known MTTKRP (matricized tensor times Khatri-Rao product) [36] operation, which is in the form of 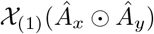, and can be computed in 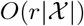 if 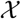 is a sparse tensor with 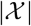 non-zeros, and *Â_x_* and *Â_y_* have *r* columns. Thus, the overall time complexity of computing Equation (3) is 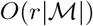.

Following the derivations in [34], we obtain the partial derivatives of the second term 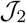 as

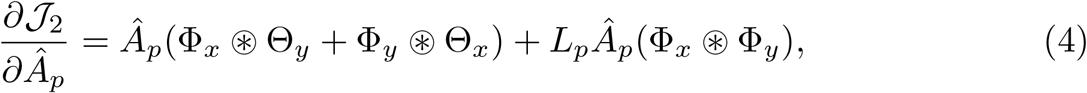

where 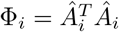, and 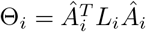, for all *i* ∈ {*x, y, p*}. It is not hard to show that the complexity of computing the Equation (4) is 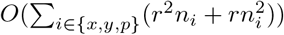.

Next, we combine 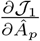 and 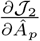 to obtain the overall derivative as

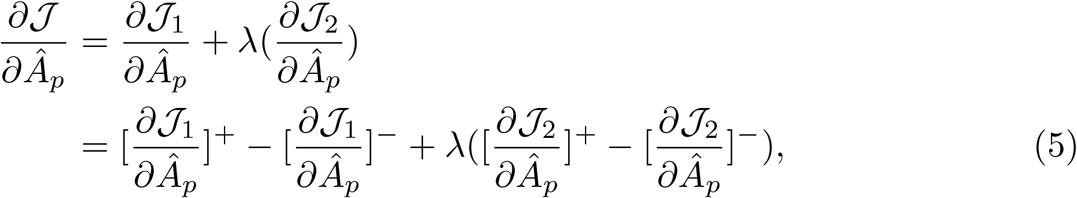

where 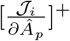 and 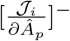 are non-negative components in 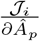, which are defined below as

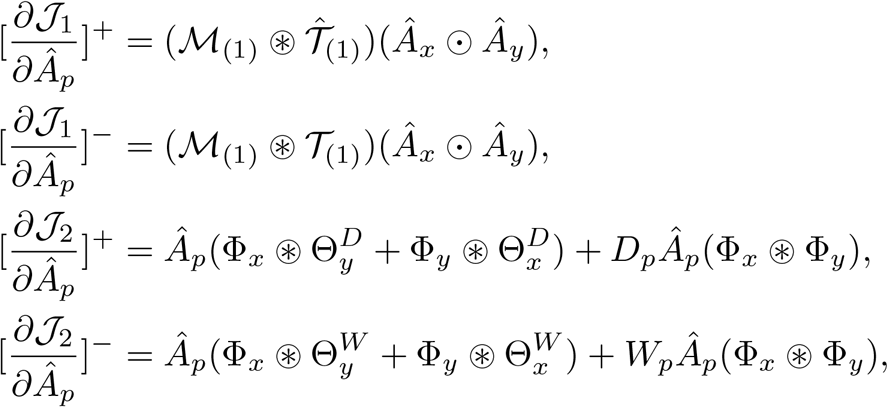

where 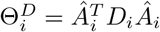 and 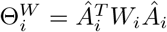, for all *i* ∈ {*x, y, p*}. According to Equation (5), the objective function 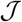 objective will monotonically decrease under the following multiplicative updating rule,

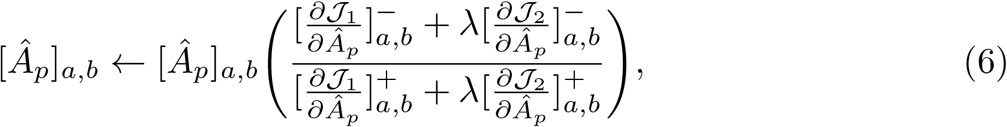

where [*Â_p_*]_*a,b*_ denotes the (*a, b*)-th element in matrix *Â_p_*. Similarly, we can derive the update rule for [*Â_x_*]_*a,b*_ and [*Â_p_*]_*a,b*_ as follows,

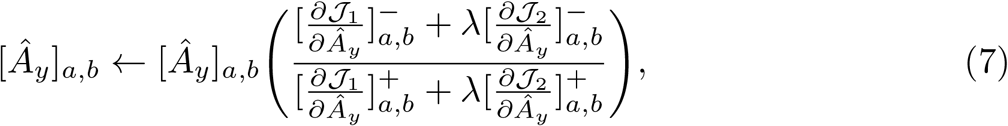

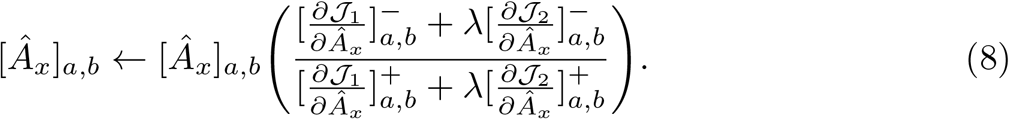

We then propose an iterative algorithm FIST in Algorithm to find the local optimum of the proposed graph regularized tensor completion problem with time complexity 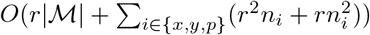. The theoretical convergence analysis of FIST is given in Appendix.

**Algorithm 1.**
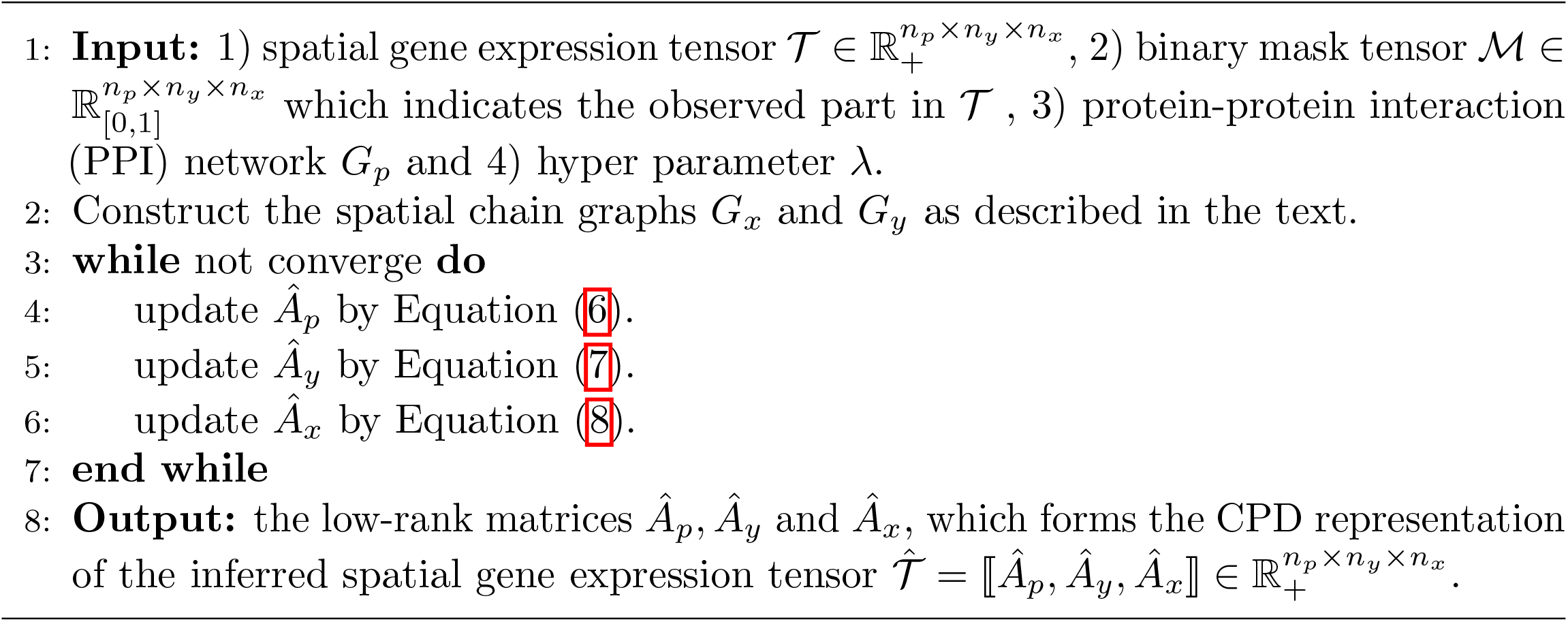
FIST: **F**ast **I**mputation of **Sp**atially-resolved Gene Expression **T**ensor

### Related methods for comparison

To benchmark the performance of FIST, we compared it with three matrix factorization (MF)-based methods (with graph regularizations) and a nearest neighbors (NN)-based method, which have been applied to impute various types of biological data including the imputation of dropouts in single-cell RNA sequencing (scRNA-seq) data. Note that NMF-based methods have been shown to be effective for learning latent features and clustering of high-dimension sparse genomic data [37].

- **ZIFA**: Zero-inflated factor analysis (ZIFA) [26] factorizes the single cell expression data 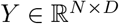 where *N* and *D* denote the number of single cells and genes respectively, into a factor loading matrix 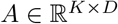 and a matrix 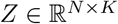 which spans the latent low-dimensional space where dropouts can happen with a probability specified by an exponential decay associated with the expression levels. The imputed matrix can be computed as *Ŷ* = *ZA* + *μ*, where 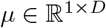 is the latent mean vector.
- **REMAP**: Since ZIFA is a probabilistic MF model which does not utilize the spatial and gene networks, we therefore also compare with REMAP [38], which was developed to impute the missing chemical-protein associations for the identification of the genome-wide off-targets of chemical compounds. REMAP factorizes the incomplete chemical-protein interactions matrix into the chemical and protein low-rank matrices, which are regularized by the compound similarity graph and protein sequence similarity (NCBI BLAST [39]) graph respectively.
- **GWNMF**: Both ZIFA and REMAP are only applicable to the spot-by-gene matrix which is a flatten of a input tensor 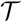. Such flattening process assumes the spots are independent from each other, and thus loses the spatial information. To keep the spatial arrangements, we also apply MF to each *n_x_* × *n_y_* slice in tensor 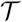. Specially, we adopt the graph regularized weighted NMF (GWNMF) [40] method to impute each *n_x_* × *n_y_* gene slice. We let GWNMF use the same *x*-axis and *y*-axis graphs *G_x_* and *G_y_* as described in the previous section to regularize the MF.
- **Spatial-NN**: It has been observed that in sparse high-dimensional scRNA-seq data, constructing a nearest neighbor (NN) graph among cells can produce more robust clusters in the presence of dropouts because of taking into account the surrounding neighbor cells [41]. Such rationale has be considered in the clustering methods such as Seurat [42] and shared nearest neighbors (SNN)-Cliq [41], and can also be adopted to impute the spatial gene expression data. We introduce a SNN-based baseline Spatial-NN using neighbor averaging to compare with FIST. Specifically, to impute the missing expression of a target spot, Spatial-NN first searches its spatially nearest spots with observed gene expressions, then assign their average gene expression to the target spot.

We used the provided Python package^1^ to experiment with ZIFA, and the provided MATLAB package^2^ to experiment with REMAP. To apply both methods, we rearranged the data tensor 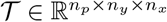 to a matrix 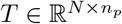, where *N* = *n_x_n_y_* denotes the total number of spots. The spatial graph of REMAP is constructed by connecting two spots if they are spatially adjacent. REMAP adopts the same PPI network as the gene graph *G_p_* as used by FIST. We used MATLAB to implement GWNMF and Spatial-NN ourselves to impute each gene slice 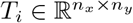 in 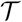. In the comparisons, the graph hyperparameter λ of FIST is only selected from {0, 0.01, 0.1, 1}. The graph hyperparameters of REMAP and GWNMF are set by searching the grids from {0.1, 0.5, 0.9, 1} and {0, 0.1, 1, 10, 100} respectively as suggested in the original studies. Note that different methods use different scales of graph hyperparameters since the gradients of their variables with respective to the regularization terms are in different scales. The optimal hyper-parameters are selected by the validation set for FIST. For FIST, REMAP and GWNMF, we applied PCA on matrix 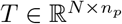 to determine the rank *r* ∈ [200, 300] of the low-rank factor matrices, such that at least 60% of the variance is accounted for by the top-*r* PCA components of *T*. The latent dimension *K* of ZIFA is set to 10 since it is time consuming to run ZIFA with a larger *K*. We also observed that increasing K from 10 to 50 does not show clear improvement on the imputation accuracy.

## Results

In this section, we first describe data preparation and performance measure and then show the results of spatial gene expression imputation. We also analyzed the results by the gene-wise density of the gene expressions and regularization by permuted graphs. Finally, we analyzed the imputed spatial gene expressions in the Mouse Kidney Section dataset to show several interesting gene clusters revealing functional characteristics of the three tissue regions, corex, OSOM and ISOM.

### Preparation of spatial gene expression datasets

We downloaded the spatial transcriptomic datasets from 10x Genomics^3^, which is a collection of spatial gene expressions in 10 different tissue sections from mouse brain, mouse kidney, human breast cancer, human heart and human lymph node as listed in Table 2. All the sptRNA-seq datasets were collected with 10x Genomics Visium Spatial protocol (v1 chemistry) [14] to profiles each tissue section with a high density hexagonal array with 4,992 spots to achieve a resolution of 55 μm (1-10 cells per spot). To fit a tensor model on the spatial gene expression datasets, we organized each of the 10 datasets into a 3-way tensor 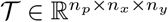, where the (*i, j, k*)-th entry in 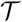 is the UMI count of the *i*-th gene at the (*k, j*)-th coordinate in the array. We set the entries in 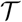 to zeros if their UMI counts is lower than 3. We then removed the genes with no UMI counts across the spots, and removed the empty spots in the boundaries of the four sides in the H&E staining from 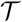. The sizes and densities of the 10 different spatial gene expression tensors after prepossessing are also summarized in Table 2. The log-transformation is finally applied to every entry of 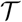 as 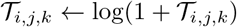. Finally, we downloaded the Homo sapiens protein-protein interactions (PPI) network from BioGRID 3.5 [31] as the gene network *G_p_* to match with the genes in each dataset.

**Table 2.** 10x Genomics spatial transcriptome data from 10 tissue sections.

| Dataset | Tissue section | Tensor dimensions | Density |
| --- | --- | --- | --- |
| HBA1 | Human Breast Cancer (Block A Section 1) | $13,426 \times 60 \times 77$ | 0.093 |
| HBA2 | Human Breast Cancer (Block A Section 2) | $13,470 \times 58 \times 75$ | 0.100 |
| HH | Human Heart | $7,487 \times 63 \times 70$ | 0.049 |
| HLN | Human Lymph Node | $12,368 \times 61 \times 78$ | 0.088 |
| MKC | Mouse Kidney Section (Coronal) | $12,264 \times 41 \times 56$ | 0.103 |
| MBC | Mouse Brain Section (Coronal) | $13,570 \times 49 \times 74$ | 0.110 |
| MB1P | Mouse Brain Serial Section 1 (Sagittal-Posterior) | $15,404 \times 62 \times 67$ | 0.115 |
| MB2P | Mouse Brain Serial Section 2 (Sagittal-Posterior) | $12,497 \times 63 \times 65$ | 0.077 |
| MB1A | Mouse Brain Serial Section 1 (Sagittal-Anterior) | $12,658 \times 59 \times 66$ | 0.105 |
| MB2A | Mouse Brain Serial Section 2 (Sagittal-Anterior) | $12,295 \times 63 \times 66$ | 0.082 |

### Performance measures

We applied 5-fold cross-validation to evaluate the performance of imputing spatial gene expressions. Specifically, for each gene, we chose 4-fold of its observed expression values (non-zeros in 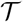) for training and validation, and hold out the rest 1-fold observed expression values as test data. The hyper-parameter λ is optimized by the validation set for FIST. Denoting vectors 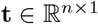 and 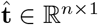 as the true and predicted expressions of a target gene on the *n* hold-out test spots, the prediction performance of the target gene is evaluated by the following three metrics,

- MAE (mean absolute error) 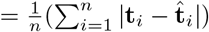,
- MAPE (mean absolute percentage error) 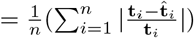,
- R^2^ (coefficient of determination) 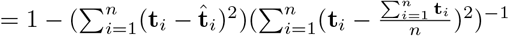.

We expect a method to achieve smaller MAE and MAPE and larger R^2^ for better performance.

### FIST signficantly improves the accuracy of imputing spatial gene expressions

The performance of gene expression imputation in the five fold cross-validation on the ten sptRNA-seq datasets are shown in Figure 2. The average and standard deviation of the prediction performances across all the genes are shown as error bar plots in Figure 2. FIST consistently outperforms all the baselines with significantly lower MAE and MAPE, and larger R^2^ in all the 10 datasets. FIST also shows a more robust performance across all the genes as the variances in all the three evaluation metrics are also lower than the other compared methods. To examine the prediction performance more closely, Figure 3 shows the distributions of MAE of individual genes in the 10 datasets (the box plots of MAPE and R^2^ are given in S1 Figure and S2 Figure). The result is consistent with the overall performance in Figure 2. The observations suggest that FIST indeed performs better than the other methods in the imputation accuracy informed by the spatial information in the tensor model. It is also noteworthy that GNMF, the MF method regularized by the spatial graph applied to each individual gene slice in tensor 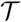, outperforms the other baselines in almost all the datasets. This observation further confirms that the spatial patterns maintained in each gene slice is informative for the imputation task. It is clear that FIST outperformed GNMF with better use of the spatial information coupled with the functional modules of the PPI network *G_p_* and the joint imputation of all the genes in the tensor 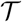.

**Fig 2.**
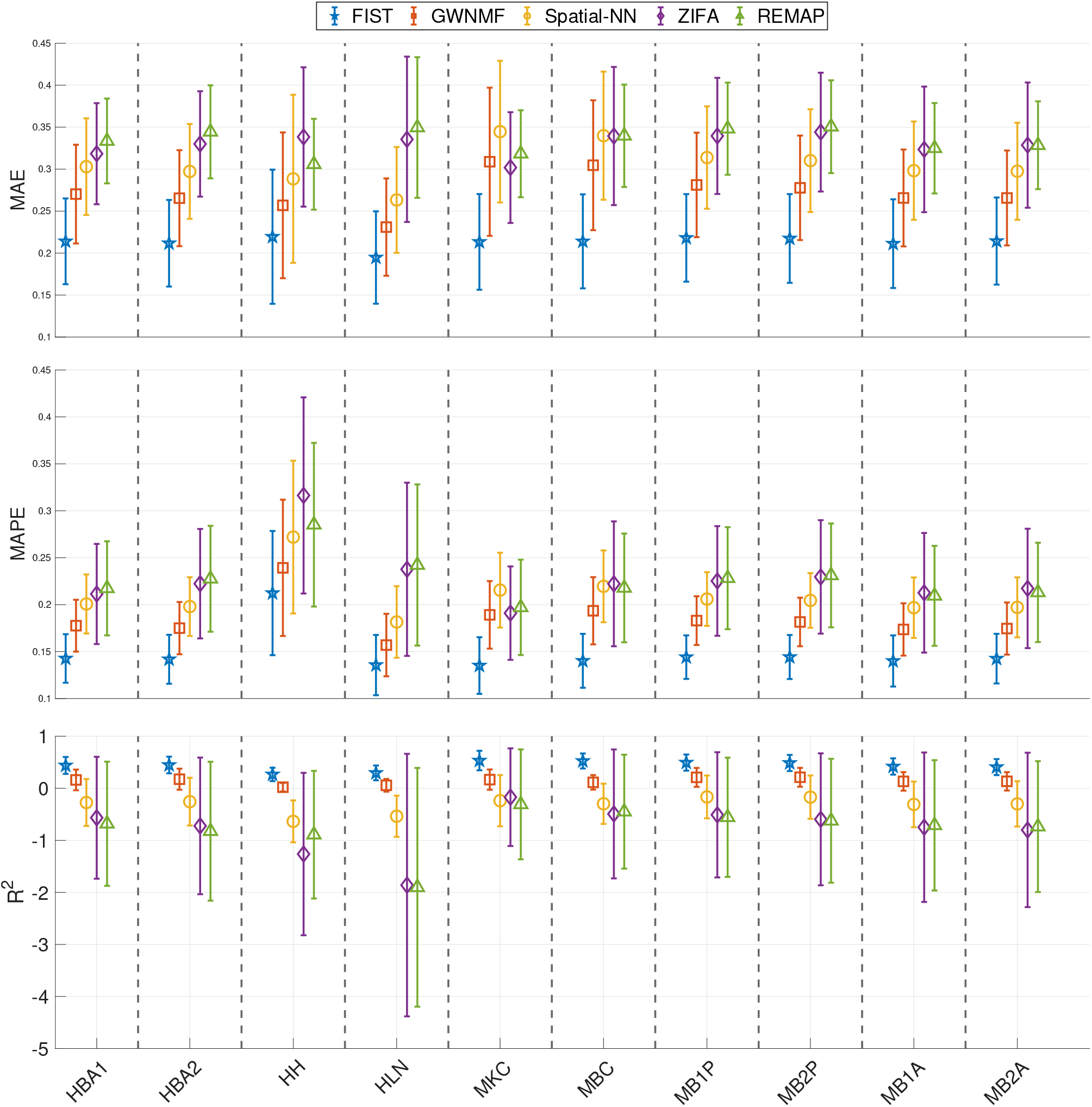
Cross-validation results on gene imputation. The performances of the five compared methods on the 10 tissue sections are measured by 5-fold cross-validation. The error bars (mean and variance) in different shapes and colors show the imputation performance of the methods on all the genes. The result on each of the 10 datasets is shown in one vertical column separated by dashed lines.

**Fig 3.**
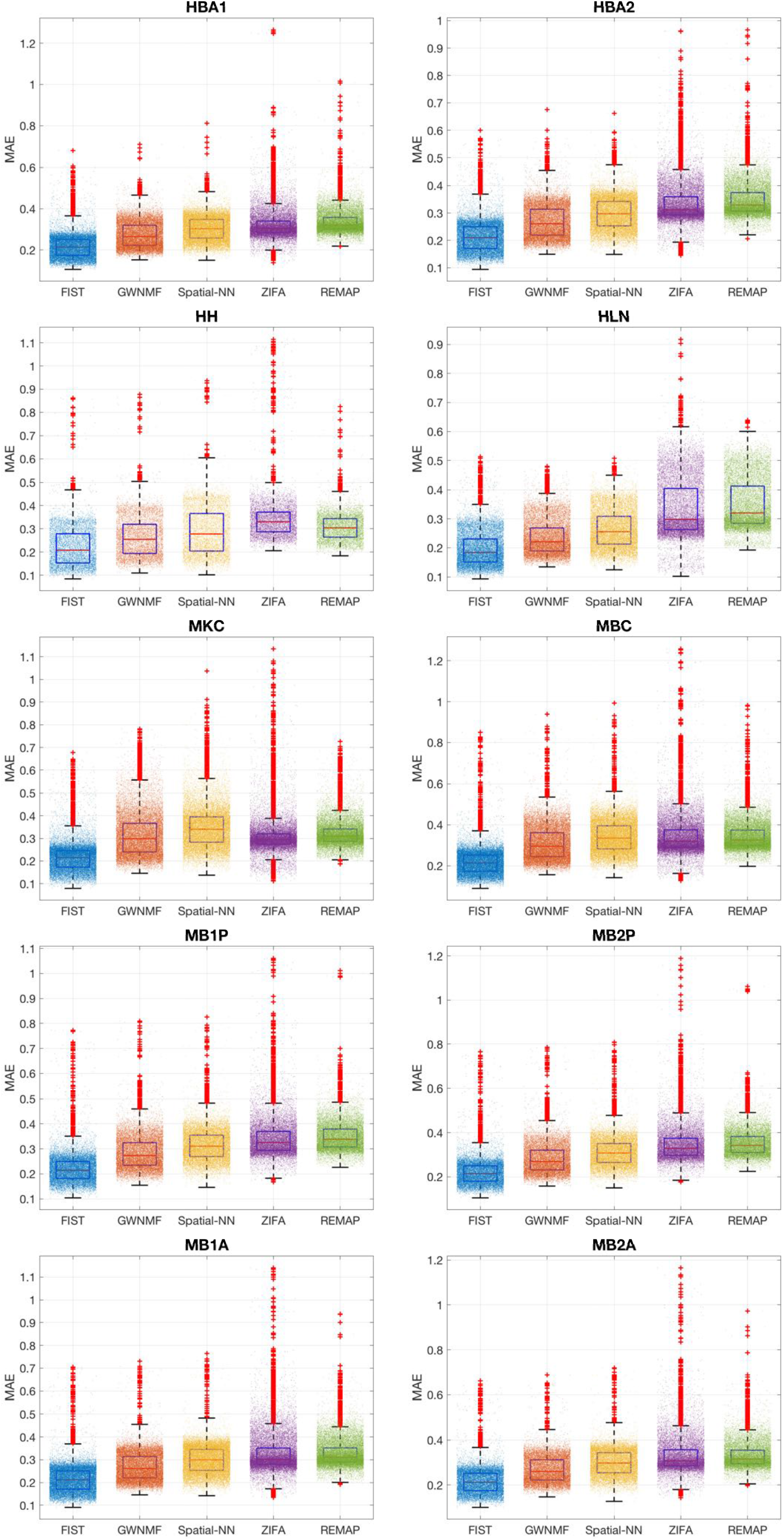
Gene-wise imputation performance by MAE. The performances on the imputations of each gene are shown as box plots. The MAE of every gene slice is denoted by one dot. The performance of each method is shown in each colored box plot.

### Cartesian product graph regularization plays a significant role

To understand the role of the regularization by the Cartesian product graph, we further analyzed the genes that are benefitting most from the regularization by the Cartesian product graph in the cross-validation experiments. In particular, in the grid search of the optimal λ weight on the CPG regularization term by the validation set, we count the percentage of the genes with optimal λ = 0.01 rather than 0, which means completely ignore the regularization. To correlate the improved imputations with the sparsity of the gene expressions, we divided all the genes into 4 equally partitioned groups (L1-4) ordered by their densities in the sptRNA-seq data, where L1 and L4 contain the sparsest and the densest gene slices, respectively. For each of the four density levels, we count the percentage of gene slices that benefit from the CPG regularization and plot the results in Figure 4(A). In the plots, there is a clear trend that the sparser a gene slice, the more likely it benefits from the CPG regularization in all the ten datasets. In the densest L4 group, as low as 20% of the genes can benefit from the CPG regularization versus more than 50% in the sparest L1 group. This is understandable that there is less training information available for sparsely expressed genes (with more dropouts) and the spatial and functional information in the CPG can play a more important role in the imputation by seeking information from the gene’s spatial neighbors or the functional neighbors in the PPI network. This observation is also consistent with the fact that the performance of tensor completion tends to degrade severely when only a very small fraction of entries are observed [43, 44], and therefore those sparser gene slices tend to benefit more from the side information carried in the CPG.

**Fig 4.**
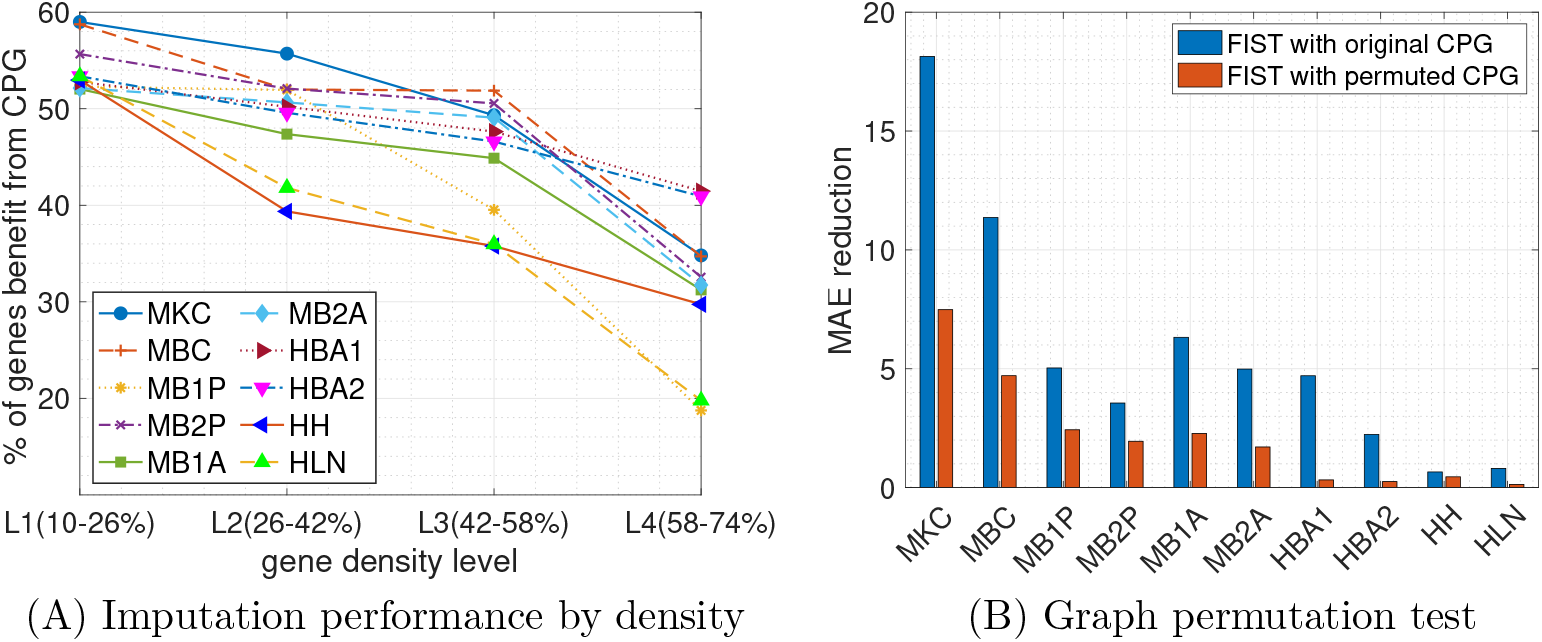
Analysis of Cartesian product graph regularization. (A) The percentages of genes benefit from the CPG are plotted by their densities in four different ranges. Each colored line represents one of the 10 datasets. (B) The total reduction of MAE using the original and permuted graphs are compared across the 10 tissue sections.

To further confirm the role of the Cartesian product graph, we also compared the performance of FIST using the CPG of *G_x_, G_y_* and *G_p_* with the one using a randomly permuted graph from the CPG. To generate the random graph, we first generated three random graphs by permute *G_x_, G_y_* and *G_p_* individually which also preserves the degree distributions of the original graphs, by randomly swapping the edges in each graph while keeping the degree of each node. Then we measured the performances of FIST using the original CPG and the CPG obtained from the permuted graphs by MAE reduction, which is the total reduction of MAE on all the genes by varying hyperparameter λ from 0 to 0.01 meaning not using the graph versus using the graph. The comparisons across all the 10 datasets are shown in Figure 4(B). We observe that the FIST using the original graphs receives much higher MAE reduction than the FIST using the permuted graphs. This observation suggests the topology in the original graph topology carry rich information that is helpful for the imputation task beyond just the degree distributions preserved in the random graphs.

### FIST imputations recover spatial patterns enriched by highly relevant functional terms

To demonstrate that imputations by FIST can reveal spatial gene expression patterns with highly relevant functional characteristics among the genes in the spatial region, we performed comparative GO enrichment analysis of gene clusters detected with the imputed gene expressions. We conducted a case study on the Mouse Kidney Section data to further analyze the associations between the spatial gene clusters and the relevance between their functional characteristics and three kidney tissue regions, corex, outer stripe of the outer medulla (OSOM) and inner stripe of the outer medulla (ISOM).

To validate the hypothesis that the imputed sptRNA-seq tensor 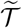 given below

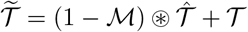

can better capture gene functional modules than the sparse sptRNA-seq tensor 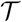 does, we first rearranged both sptRNA-seq tensors into matrices 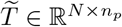 and 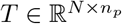, where *N* = *n_x_n_y_* denotes the total number of spots. We then applied K-means on each matrix to partition the genes into 100 clusters. Next, we used the enrichGO function in the R package clusterProfiler [45] to perform the GO enrichment analysis of the gene clusters. The total number of significantly enriched gene clusters (FDR adjusted p-value < 0.05) in each of the 10 tissue sections are shown in Figure 5, which clearly tells that K-means on the imputed sptRNA-seq data produces much more significantly enriched clusters across all the 10 tissue sections than the sparse sptRNA-seq data without imputation.

**Fig 5.**
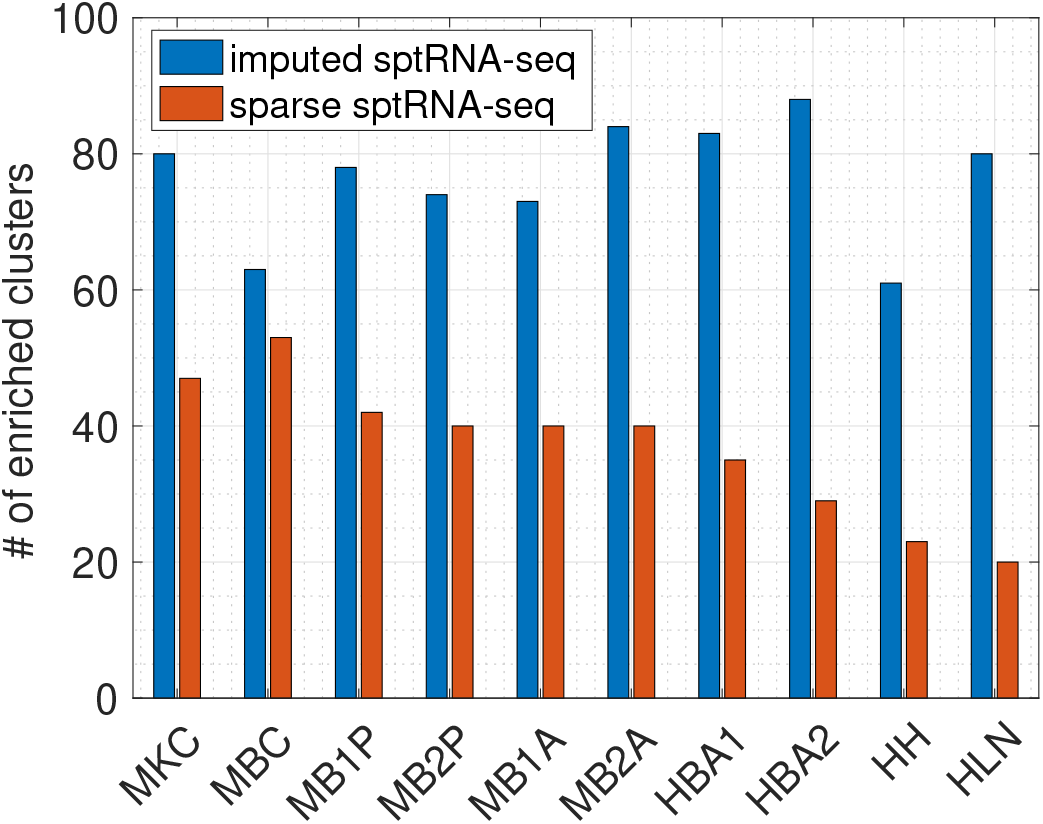
Enrichment analysis on the sparse and imputed sptRNA-seq data. The total number of significantly enriched clusters (with at least one enriched GO term with FDR adjusted p-value < 0.05) in the 10 tissue sections are shown.

Finally, we conducted a case study on the Mouse Kidney Section and present the highly relevant functional characteristics in different tissues in mouse kidney detected with the imputations by FIST. For each of the 100 gene clusters generated by K-means as described above, we collapsed the corresponding gene slices in 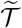 into a *n_x_* × *n_y_* matrix by averaging the slices to visualize the center of the gene cluster. The visualizations and enrichment results of all the 100 clusters are given in Table S1. We focus on 3 kinds of representative clusters in Figure 6 which match well with three distinct mouse kidney tissue regions: cortex, ISOM (inner stripe of outer medulla and OSOM (outer stripe of outer medulla). By investigating the enriched GO terms by the clusters (*p*-values shown in Table 3), we found their functional relevance to cortex, ISOM and OSOM regions. We found that the spatial gene cluster 9 which is highly expressed in cortex specifically enriched biological processes for the regulation of blood pressure (GO:0008217, GO:0003073, GO:0008015 and GO:0045777) and transport/homeostasis of inorganic molecules (GO:0055067 and GO:0015672). The spatial gene cluster 23 and 28 which are also highly expressed in cortex enriched cellular pathways that are critical for the polarity of cellular membranes (GO:0086011, GO:0034763, GO:1901017, GO:0032413 and GO:1901380) and the transport of cellular metabolites (GO:1901605, GO:0006520, GO:0006790 and GO:0043648), respectively. These observations are consistent with previous studies reporting the regulation of kidney function by above listed biological processes in cortex [46–49]. In contrast, the pattern analysis of spatial gene expression in cluster 4, 8, 25 and 52 which are highly expressed in OSOM in kidney showed that catabolic processes of organic and inorganic molecules are specifically enriched such as GO:0015711, GO:0046942, GO:0015849, GO:0015718, GO:0010498, GO:0043161, GO:0044282, GO:0016054, GO:0046395, GO:0006631, GO:0072329, GO:0009062 and GO:0044242. These cellular processes are known to be active in renal proximal tubule which exists across cortex and OSOM [50–55]. Distinctively, the spatial gene clusters highly expressed in ISOM enriched pathways for nucleotide metabolisms (GO:0009150, GO:0009259 and GO:0006163) in cluster 3 and renal filtration (GO:0097205 and GO:0003094) in cluster 5. Collectively, these observations demonstrate that FIST could identify physiologically relevant distinctive spatial gene expression patterns in the mouse kidney dataset. Further, it suggests that FIST can provide a high-resolution anatomical analysis of organ functions in sptRNA-seq data.

**Fig 6.**
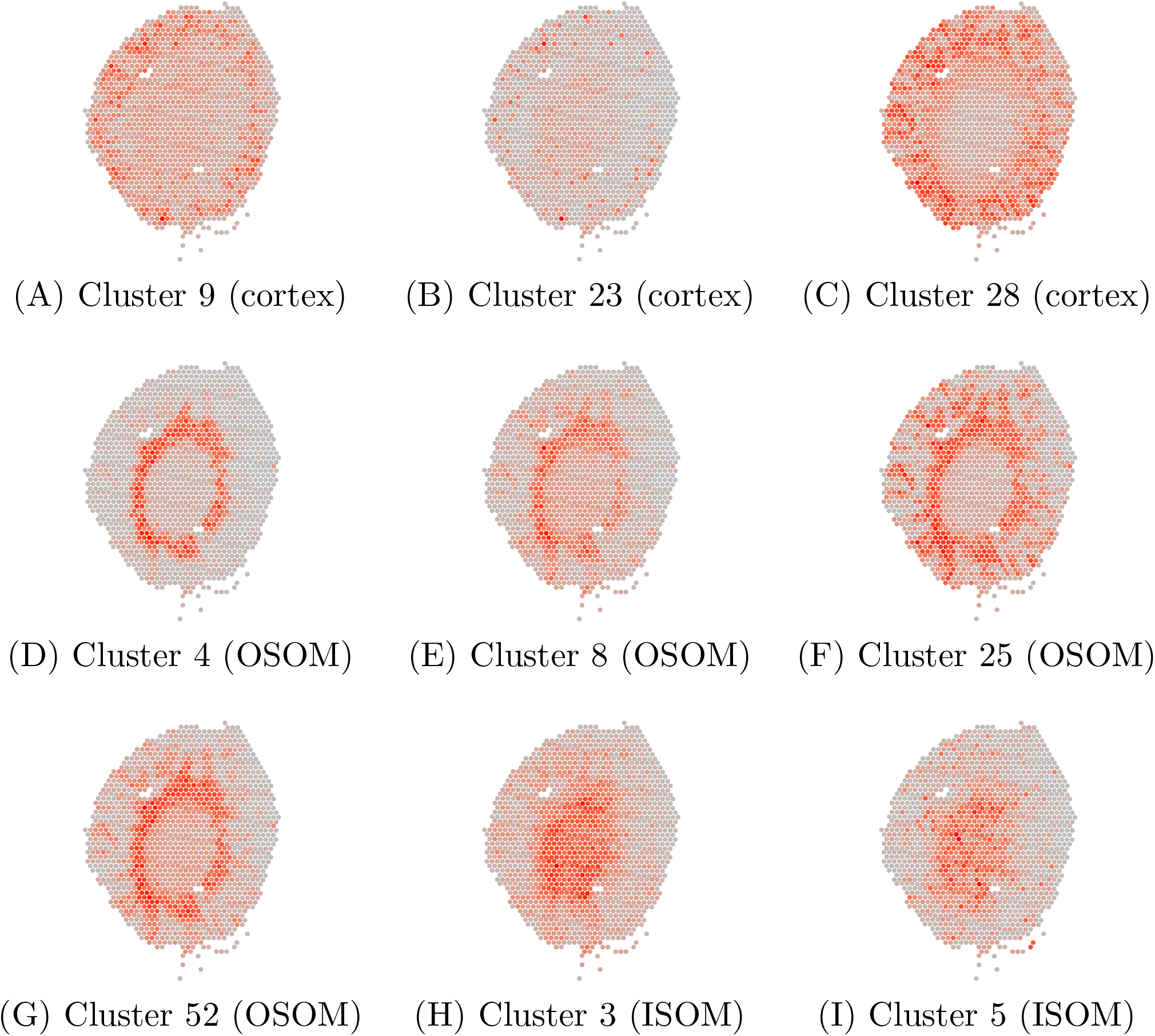
FIST recovers spatial gene expression patterns on Mouse Kidney Section. (A)-(C) Gene expression patterns active in the cortex region, (D)-(G) Gene expression patterns active in the outer stripe of the outer medulla (OSOM) region, (H)-(I): Gene expression patterns active in the inner stripe of the outer medulla (ISOM) region.

**Table 3.**
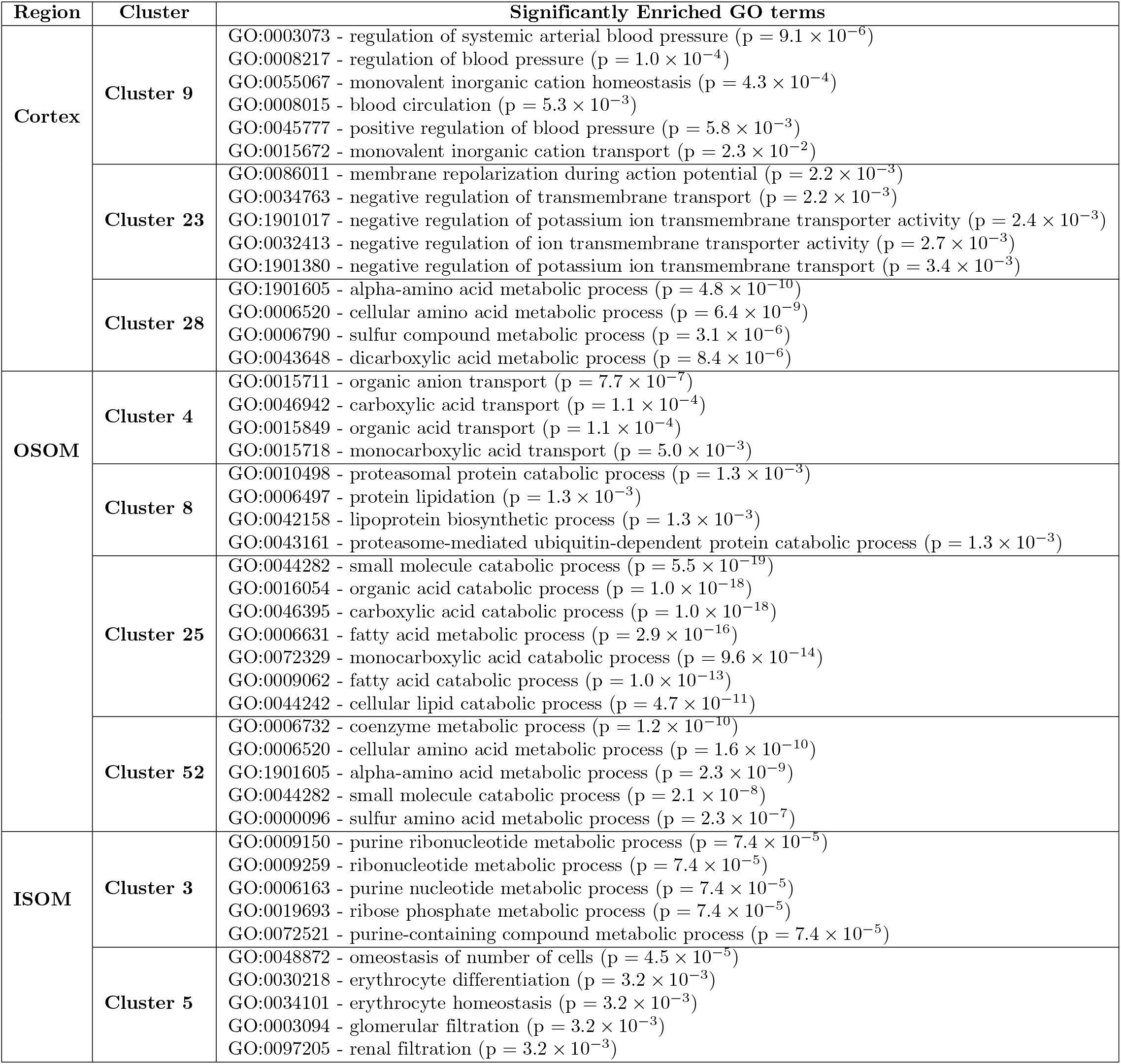
Functional terms enriched by spatial gene clusters (most significant relevant functions)

## Discussions

In this study, we proved that tensor is a natural representation of the multidimensional structure in spatially-resolved gene expression data mapped by the 2D spatial array. To the best of our knowledge, this is the first work to model the imputation of spatially-resolved transcriptomes as a tensor completion problem. Our key observations in the experiments with the ten 10x Genomics Visium spatial transcriptomic datasets are that 1) the imputation accuracy is significantly improved by leveraging the tensor representation of the sptRNA-seq data, and 2) by incorporating the spatial graph and PPI network, the accuracy the imputation and the content of the functional information in the imputed spatial gene expressions can be further improved significantly.

We observed that the genes that are more sparsely expressed can benefit more from the adjacency information in the spatial graph and the functional information in the PPI network. These genes can be empirically detected with a validation set to tune the only hyper-parameter λ for deciding if the regularization by the product graph is needed for the imputation of a gene. Thus, we expect a low risk of overfitting in applying FIST to other datasets. In addition, the functional analysis of the spatial gene clusters detected on the Mouse Kidney Section data further confirms that FIST detects gene clusters with more spatial characteristics that are consistent with the physiological features of the tissue.

Although our experiments focused on medium density 10x Genomics Visium kit array (5000 spots), we also further tested that FIST is also applicable and scalable to high-resolution spatial transcriptomics datasets with millions of spots in the preliminary work in a follow-up study. We tested the high-definition spatial transcriptomics (HDST) datasets generated from [14]. The HDST datasets includes 3 mouse tissue sections from olfactory bulb and 3 human tissue sections from breast cancer using hexagonal array to profile tissue with a high density (1,467,270 spots in total) to achieve a resolution of 2 μm. Our preliminary result suggest that FIST can finish imputing each of the the 6 HDST datasets in v 1hr.

Overall, we concluded that FIST is an effective and easy-to-use approach for reliable imputation of spatially-resolved gene expressions by modeling the spatial relation among the spots in the spatial array and the functional relation among the genes. The imputation results by FIST is both more accurate and functionally interpretable. FIST is also highly generalizable to other spatial transcriotomics datasets with high scalability and only one hyper-parameter needed to tune.

## Supporting information

**S1 Figure.**
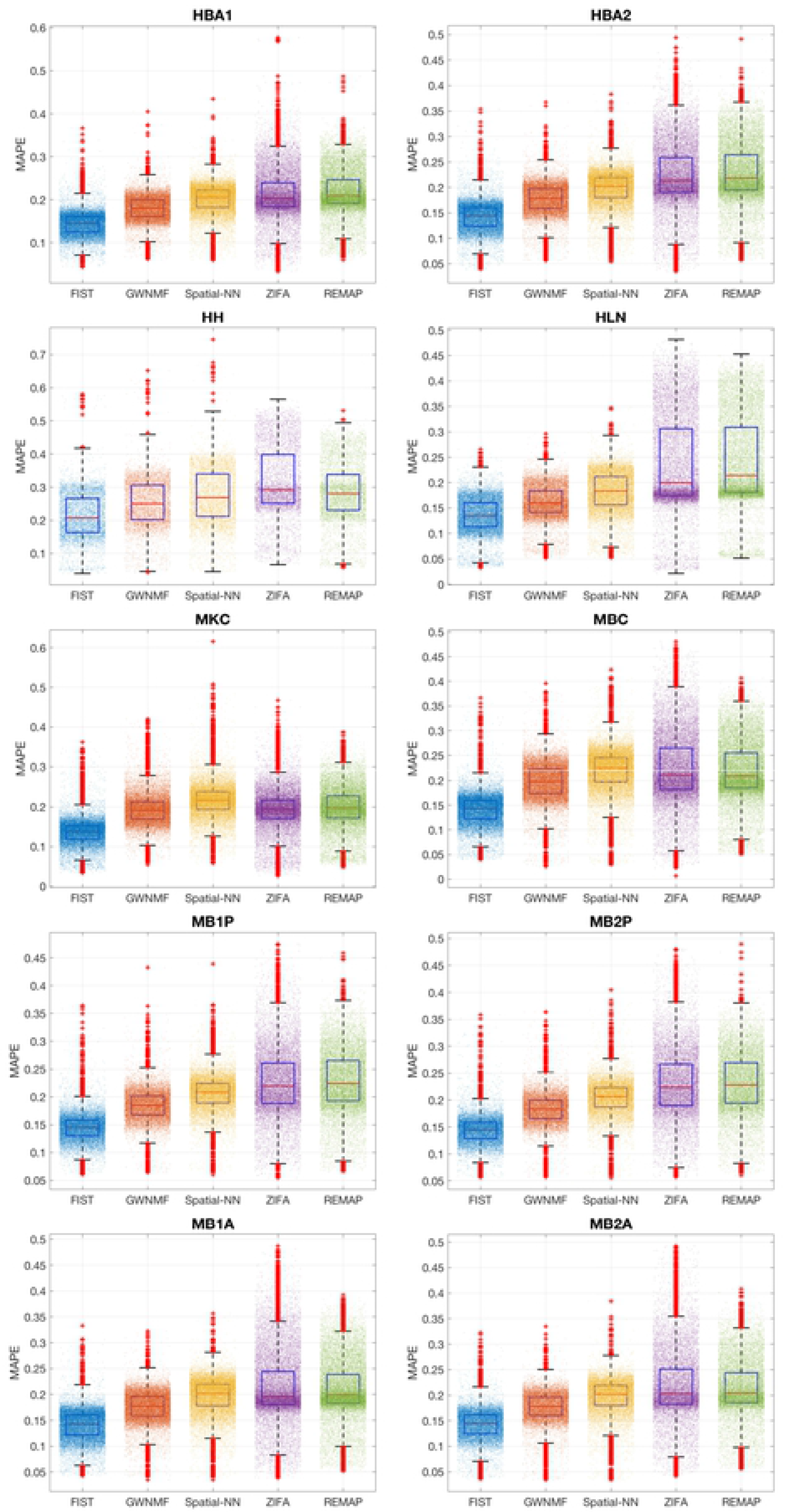
Gene-wise imputation performance by MAPE. The performances on the imputations of each gene are shown as box plots. The MAPE of every gene slice is denoted by one dot. The performance of each method is shown in each colored box plot.

**S2 Figure.**
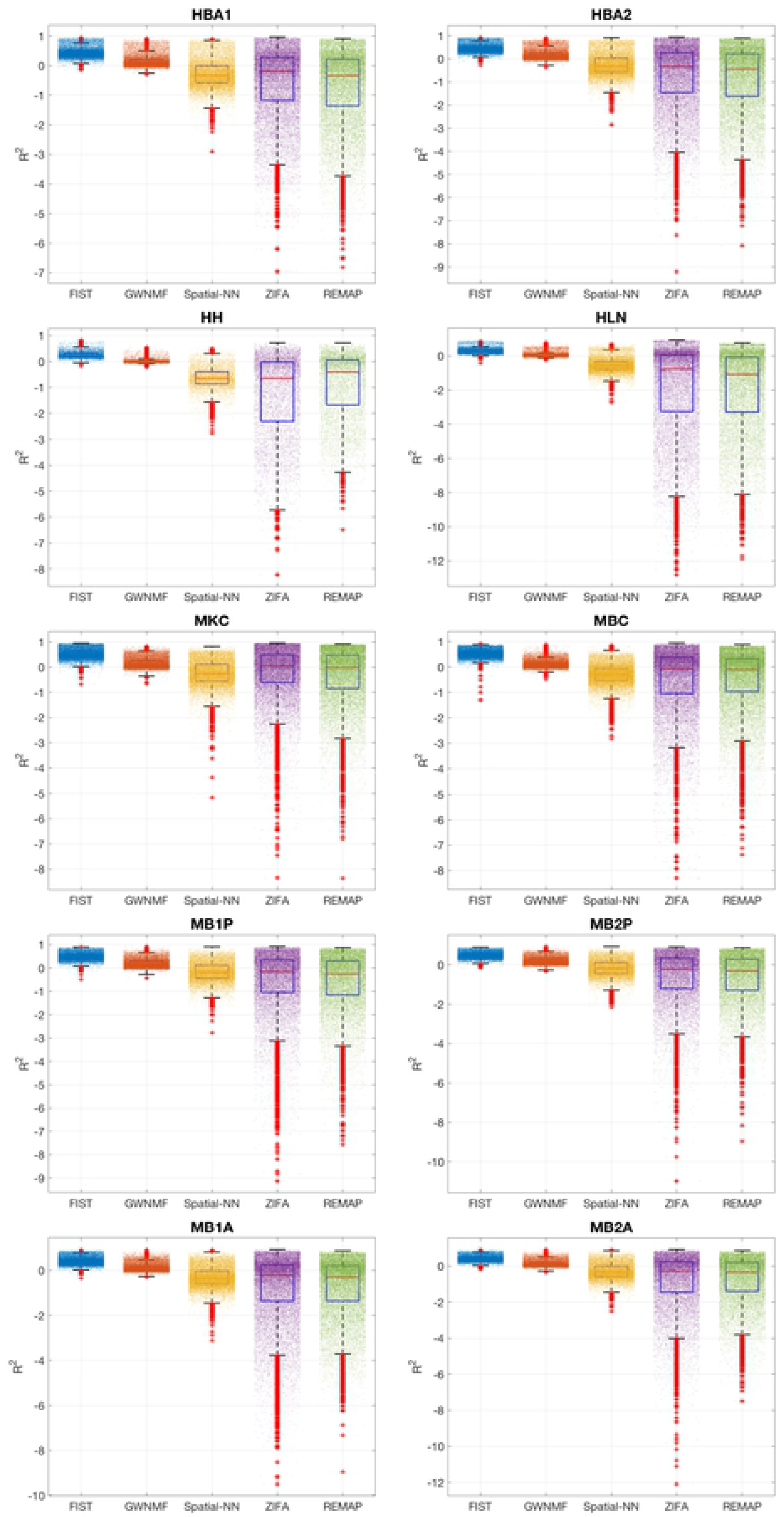
Gene-wise imputation performance by R^2^. The performances on the imputations of each gene are shown as box plots. The R^2^ of every gene slice is denoted by one dot. The performance of each method is shown in each colored box plot.

**S1 Table Enriched GO terms of spatial gene clusters.** The GO terms significantly enriched by the genes in each spatial gene cluster (FDR adjusted p-value < 0.05) are shown in the spreadsheet tables.

## S1 Appendix. Convergence of FIST

We follow the convergence analysis in our previous work [34] to show that FIST can converge under the updating rules in Equation (6)-(8).

As the objective function 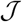 in Equation (2) is bounded from below by zero, we can prove the convergence of FIST by showing that 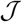 is non-increasing under each of the updating rules in Equations (6)-(8). Here, we only show that 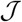 is non-increasing under Equations (6). The proof is directly applicable to Equations (7) and (8).

We first expand the derivative in Equation (5) as

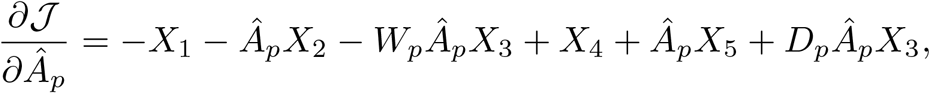

where 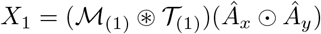, 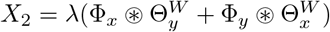, 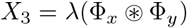, 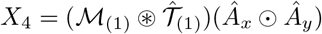, and 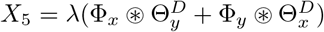.

### Theorem 1.

*Lee and Seung [35]: A function* 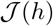 *is non-increasing under the update* 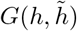 *if G*(*h, h*) *is an auxiliary function for* 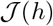, such that the following conditions are satisfied:

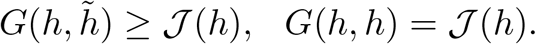

Based on Theorem 1, 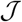 is non-increasing under the update in Equation (6) if it is an update of one proper *auxiliary function* of 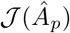, which is defined in Theorem 2.

### Theorem 2.

*The following function*

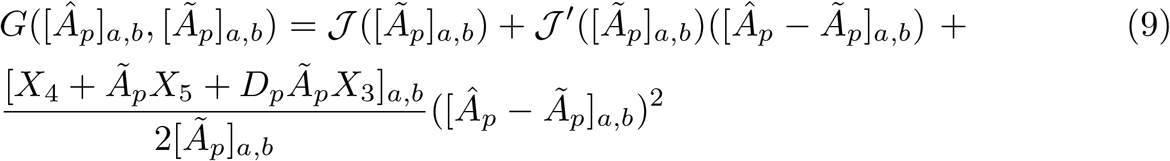

*is an auxiliary function of J* 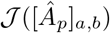 *and has its global minimum*.

Proof: First, it is obvious that 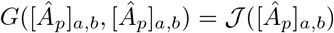. To show 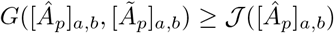 we obtain the second-order Taylor expansion of 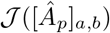 at the point [*Ã_p_*]_*a,b*_ as

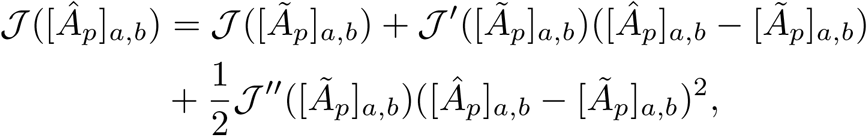

with the second-order derivative given below:

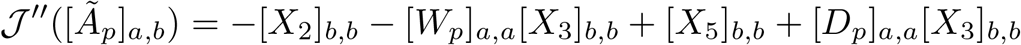

Thus, the inequality 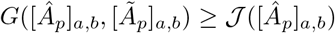 holds if

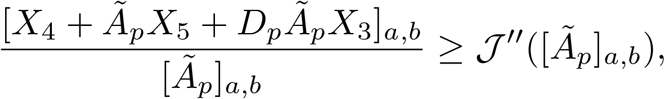

which can be demonstrated by the facts that

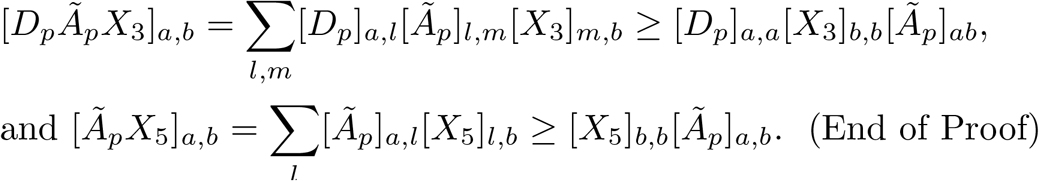

As the *auxiliary function G*([*Â_p_*]_*a,b*_, *Â_p_*]_*a,b*_) in Equation (9) is a quadratic function on variable [*Â_p_*]_*a,b*_, its minimum can be easily obtained in a closed-form as

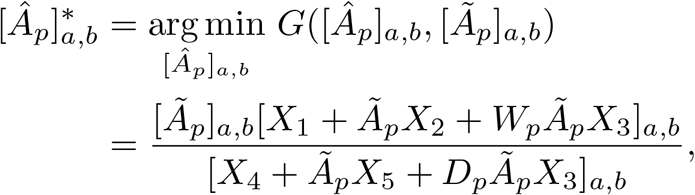

which leads to the updating rule in Equation (6).

To analyze the optimality of the fixed point after convergence, we first define 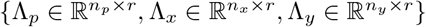 to be the matrices of Lagrange multipliers with the Lagrange function

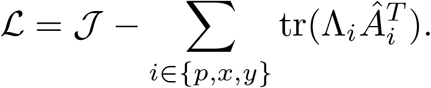

Setting 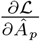 to be zero, we obtain 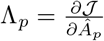. Furthermore, when *A*^(*i*)^ is a fixed point under the updating in Equation (6) we have

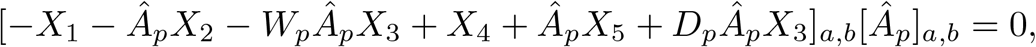

which implies the KKT complementary slackness condition [Λ_*p*_]_*a,b*_ [*A_p_*]_*ab*_ = 0 is satisfied.

## Acknowledgments

This research work was supported by University of Minnesota - Minnesota Robotics Institute Seed Grants and NIH grant R01GM113952-01A1. The funders had no role in study design, data collection and analysis, decision to publish, or preparation of the manuscript.

## Footnotes

1 https://github.com/epierson9/ZIFA

2 https://github.com/hansaimlim/REMAP

3 https://support.10xgenomics.com/spatial-gene-expression/datasets/

